# Antibiotic Exposure Through Human Milk Influences the Infant Gut Microbiome

**DOI:** 10.64898/2026.03.31.715750

**Authors:** Kine Eide Kvitne, Simone Zuffa, Vincent Charron-Lamoureux, Abubaker Patan, Julius Agongo, Jiaxi Cai, Victoria Deleray, Yasin El Abiead, Shipei Xing, Jasmine Zemlin, Sydney Thomas, Megan R. Nelson, Abigail Gant, Angelique Ghadishah, Anny Lam, Benjamin Ho, Jeremiah D. Momper, Raymond T. Suhandynata, Kerri Bertrand, Rob Knight, Christina Chambers, Pieter C. Dorrestein, Shirley M. Tsunoda

**Affiliations:** Skaggs School of Pharmacy and Pharmaceutical Sciences, University of California San Diego, La Jolla, CA, USA; Shu Chien-Gene Lay Department of Bioengineering, University of California San Diego, La Jolla, USA; Department of Pathology, University of California San Diego, La Jolla, USA; Department of Pediatrics, University of California San Diego, La Jolla, USA; Center for Microbiome Innovation, University of California San Diego, La Jolla, CA, USA; Department of Computer Science and Engineering, University of California San Diego, La Jolla, CA, USA; Shu Chien-Gene Lay Department of Bioengineering, University of California San Diego, La Jolla, CA, USA; Halıcıoğlu Data Science Institute, University of California San Diego, La Jolla, CA, USA; Hong Kong University of Science and Technology Jockey Club Institute for Advanced Study, Hong Kong University of Science and Technology, Hong Kong SAR, China

**Keywords:** antibiotics, lactation, infant, dyads, early-life, microbiome, metabolomics

## Abstract

Infant antibiotic treatment is associated with increased risk of developing non-communicable diseases, potentially through disruption of the gut microbiome. However, the impact of indirect antibiotic exposure via human milk remains largely unexplored. Here, we investigate a cohort (*n*=80) of antibiotic-treated breastfeeding mother-infant dyads and untreated matching controls using integrative multi-omics analyses of fecal, milk, and skin samples (*n*=1,455). Maternal antibiotic treatment was associated with different infant fecal microbiome and metabolome profiles, including lower abundance of *Bacteroides*, *Lactobacillus*, and *Bifidobacterium*, and higher levels of antimicrobial resistance gene reads. Further, fecal metabolic alterations associated with indirect antibiotic exposure were exacerbated by formula milk supplementation. In a subset of infants (*n*=61), indirect exposure was associated with higher body mass index (BMI). These findings suggest that maternal antibiotic treatment during lactation may influence the early-life infant gut microbiome with potential long-term implications.

## INTRODUCTION

Antibiotics are life-saving medications essential for both the treatment and prevention of bacterial infections. However, antibiotic exposure disrupts the gut microbiome composition and functionality^1^. This is particularly relevant during the first year of life, when the assembly of gut microbial communities coincides with critical developmental processes in the host, including maturation of the immune system^2,3^, neurodevelopment^4^, and metabolic programming^5,6^. Several external factors, such as delivery mode^7^ and feeding practices^8,9^, can influence the structure of these microbial communities and timing of colonization^10^, leading to both immediate and long-term repercussions on health. Antibiotic exposure also perturbs this process, reducing gut microbial diversity^11^, depleting beneficial taxa, such as *Bifidobacterium* species^12^, and facilitating the enrichment of antimicrobial resistance (AMR) genes harbored by bacteria^13,14^. Several studies have associated early-life antibiotic treatment and gut microbiome disruption with increased risk of developing non-communicable diseases later in life, such as asthma^15^, allergy^15^, and obesity^16,17^.

While the effects of direct antibiotic exposure during infancy are well established^18,19^, the consequences of indirect exposure via human milk remain poorly understood^20^. This knowledge gap stems in part from the historical exclusion of pregnant and lactating women from clinical trials. Nevertheless, antibiotics are among the most frequently prescribed medications to lactating mothers^21^, with use ranging from ∼7% in high-income countries, such as in Norway^22^ and Germany^23^, to nearly 40% in low- and middle-income countries^24^. Antibiotics are frequently prescribed to treat postpartum complications, including mastitis, endomyometritis, and common respiratory or urinary tract infections, or as prophylaxis after cesarean delivery (C-section). Following maternal administration, some antibiotics can transfer into human milk depending on their physicochemical and pharmacokinetic properties^25,26^ thereby representing a potential indirect route of low-dose exposure for breastfeeding infants.

Antibiotic concentrations in milk are typically below therapeutic infant doses. These levels are typically considered safe when the relative infant dose (RID), which expresses infant drug intake as a percentage of the maternal weight-adjusted dose, is less than 10%^27^. However, this benchmark was developed to assess the risk of direct pharmacologic or toxic effects in the infant, and may not capture potential ecological effects of antimicrobial exposure. Indeed, preclinical studies have shown that maternal exposure to antibiotics can alter gut microbiome composition and offspring behavior^28,29^, and low level antibiotic exposure can increase selective pressure on the microbial resistome *in vitro*^30^. As such, indirect antibiotic exposure via human milk may have clinically relevant, long-lasting effects on the developing infant gut microbiome.

Here, we conducted a prospective clinical study leveraging the Mommy’s Milk Human Milk Research Biorepository infrastructure^31^ at UC San Diego to characterize the impact of maternal antibiotic treatment during breastfeeding on the gut microbiome of mothers and their infants. Using an integrated approach - including multi-omics analyses, such as untargeted metabolomics and metagenomics, cross-body site sampling, and targeted quantitative measurement of antibiotics in human milk - we profiled biological samples from 80 mother-infant dyads. Antibiotic exposure was associated with shifts in both maternal and infant fecal microbial and metabolic profiles and increased abundance of AMR mapping reads compared to unexposed controls. Notably, the antibiotic fecal metabolic signature in infants appeared to be exacerbated by the use of formula milk, suggesting potential interactions between antibiotic exposure and other early-life determinants of microbiome development. Additionally, in a subset of infants (*n*=61) with anthropometric data, indirect antibiotic exposure was associated with higher body mass index (BMI), highlighting the need for further investigations into potential long-term health implications of indirect antibiotic exposure.

## RESULTS

In this study, we enrolled 80 breastfeeding mother-infant dyads (**Fig. 1a**). Of these, 40 mothers received antibiotics based on clinical indication (ABX group) and the remaining mothers served as untreated matched controls (No ABX group). Paired mother-infant fecal and skin samples, together with maternal milk, were collected at baseline and during treatment. Shotgun metagenomic sequencing was performed on fecal samples (*n*=573), while untargeted liquid chromatography-tandem mass spectrometry (LC-MS/MS) metabolomics was performed on feces (*n*=573), skin (*n*=590), and human milk (*n*=292). Additionally, targeted LC-MS was used to quantify antibiotic concentrations in milk collected from treated mothers (*n*=144).

**Fig. 1.**
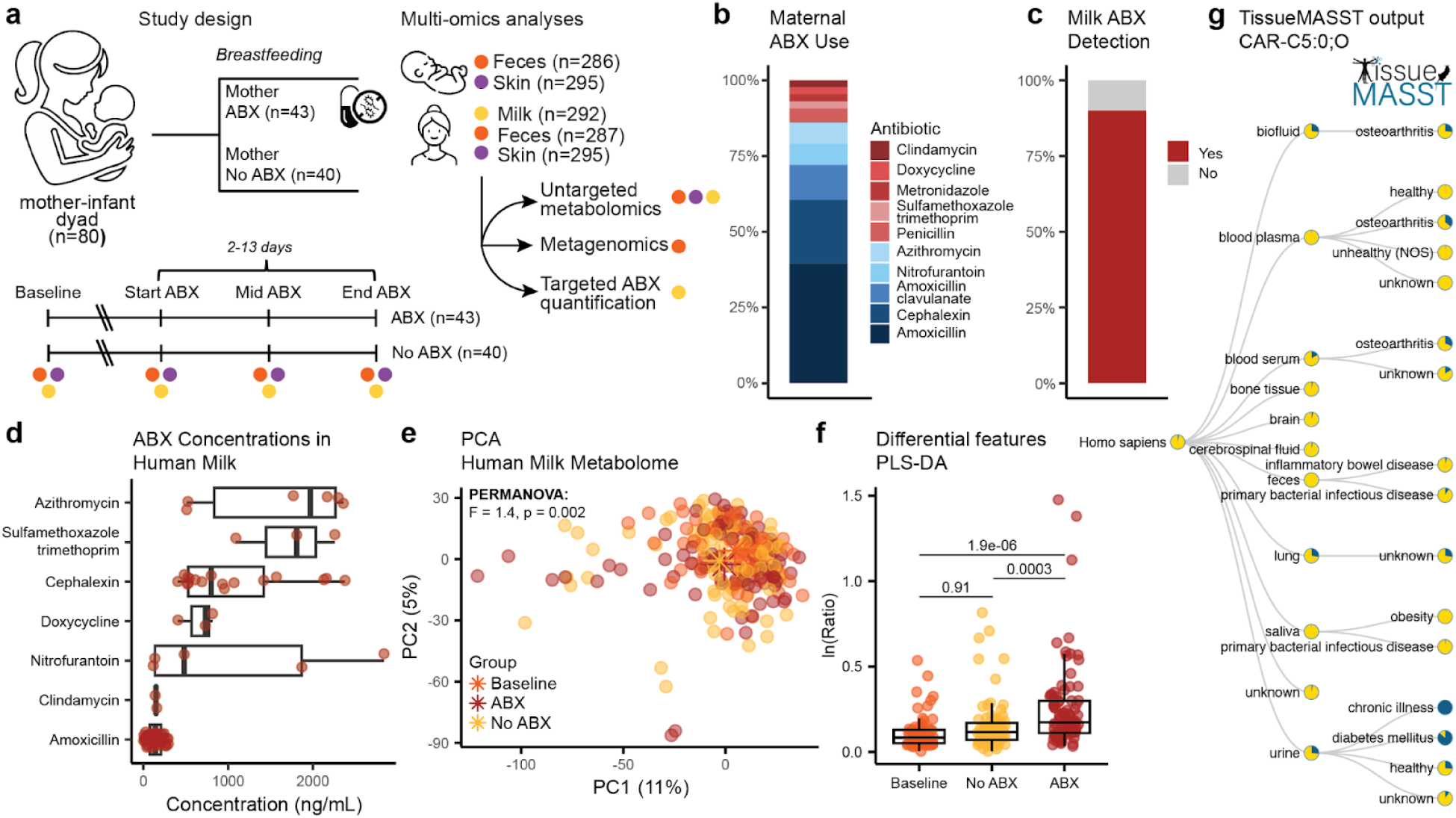
Antibiotic transfer into human milk and associated metabolic changes. **a,** Study design. Samples from breastfeeding mothers receiving antibiotics, untreated matched controls, and their infants were collected across body sites. **b,** Antibiotics used by treated mothers. **c,** Detection of antibiotics in milk samples from treated mothers using targeted LC-MS. Antibiotic detection is reported at the subject level. **d,** Boxplot of measured antibiotic concentrations in milk samples from treated mothers. Multiple samples per subject were collected. Concentrations reported for amoxicillin include samples from mothers treated with either amoxicillin or amoxicillin-clavulanate. **e**, PCA of milk metabolic profiles showing group separation (PERMANOVA; *p*=0.002). **f**, Boxplot of the natural log ratio of discriminant features enriched (numerator; top 100) or depleted (denominator; top 100) in response to ABX identified via PLS-DA. Adjusted *p* values obtained from LME with subject as random effect. **g,** TissueMASST search output for carnitine-C5:0;O. Boxplots show the first (lower), median, and third (upper) quartiles, with whiskers 1.5 times the interquartile range.

Characteristics of both mothers and infants are summarized in **Table 1**. Briefly, both groups presented comparable distributions for infant sex and delivery mode (Fisher’s Exact test, *p*>0.2). Additionally, no differences in feeding practices, methods, and frequency were observed. Infant and maternal ages were also comparable between groups (Welch’s T-test, *p*>0.45), in line with the study protocol. A tendency toward higher maternal BMI in the ABX group was observed, but it did not reach statistical significance (Welch’s T-test, *p*=0.07). Respiratory (46.5%) and urogenital infections (20.9%) were the main indications for maternal antibiotic prescription. Amoxicillin, or amoxicillin with clavulanate, were the most frequently prescribed antibiotics (51%), followed by cephalexin (21%), azithromycin (7%), and nitrofurantoin (7%) (**Fig. 1b**). Amoxicillin and cephalexin are both β-lactam antibiotics with overlapping activity against Gram-positive and some Gram-negative bacteria.

**Table 1.** Characteristics of the mother-infant dyads.

| Characteristics | ABX (n=43)<sup>†</sup> | No ABX (n=40) | P value |
| --- | --- | --- | --- |
| Infant sex, Female | 23 (53.5%) | 22 (55%) | 1 |
| Infant age (Months)<sup>#</sup> | 7.1 ± 2.91 | 7.4 ± 2.82 | 0.68 |
| Delivery mode, Vaginal | 28 (65.1%) | 31 (77.5%) | 0.24 |
| Type of delivery, Full term | 38 (88.4%) | 38 (95%) | 0.43 |
| Infant gestational age at birth (Weeks) | 38.7 ± 2.26 | 39.1 ± 1.66 | 0.26 |
| Maternal age at delivery (Years) | 33.4 ± 4.15 | 34.1 ± 4.28 | 0.45 |
| Maternal BMI (kg/m<sup>2</sup>)<sup>#</sup> | 28.3 ± 6.70 | 25.8 ± 5.32 | 0.065 |
| Household income, > \$100,000<sup>#</sup> | 31 (72.1%) | 27 (67.5%) | 0.81 |
| Ethnicity, Non-Hispanic / Non-Latina | 27 (62.8%) | 30 (75%) | 0.25 |
| Feeding practices<sup>#</sup> |  |  |  |
| - Exclusive breastmilk | 29 (67.4%) | 29 (70%) | 0.64 |
| - Mixed feeding<sup>+</sup> | 14 (32.6%) | 11 (30%) |  |
| Feeding methods<sup>#</sup> |  |  |  |
| - Breast and Bottle | 32 (%) | 25 (%) | 0.43 |
| - Breast only | 8 (%) | 10 (%) |  |
| - Bottle only | 3 (%) | 5 (%) |  |
| Feeding frequency (Times per day)<sup>#</sup> | 7.6 ± 5.11 | 7.2 ± 3.09 | 0.70 |
| Infant eating solid food, Yes<sup>#</sup> | 30 (69.8%) | 33 (82.5%) | 0.21 |
| Indication for antibiotic treatment<sup>‡</sup> |  | NA |  |
| - Respiratory infection | 20 (46.5%) |  |  |
| - Urogenital infection | 9 (20.9%) |  |  |
| - Soft tissue infection | 2 (4.7%) |  |  |
| - Mastitis | 2 (4.7%) |  |  |
| - Ear infection | 3 (7.0%) |  |  |
| - Root canal infection | 2 (4.7%) |  |  |
| - Other<sup>*</sup> | 5 (11.6%) |  |  |

<sup>†</sup>Three dyads initially enrolled as controls (No ABX) later received antibiotics and thus they were re-enrolled in the ABX group.
<sup>#</sup>At the start of antibiotic treatment in the ABX group and the corresponding timepoint in the matching controls (No ABX).
<sup>+</sup>Supplemented with formula or donor breast milk.
<sup>\*</sup>Surgery, dental surgery, C-section infection, hidradenitis suppurativa, and hysterectomy bacterial exposure.
Welch's T-test was used for continuous data. Fisher's Exact test was used for categorical data.

### Antibiotics are detectable in human milk and are associated with different milk metabolic profiles

Targeted LC-MS quantification revealed detectable antibiotic concentrations in 90% of human milk samples from treated mothers (**Fig. 1c**), suggesting that human milk may serve as a source of antibiotic exposure to breastfed infants. Concentrations varied within and between antibiotic classes, ranging from 1.6 to 2840 ng/mL **(Fig. 1d)**. Based on typical infant milk intake (up to ∼150 mL/kg/day)^32^, the measured concentrations correspond to estimated infant exposures in the μg/kg/day range, substantially below therapeutic pediatric doses for most antibiotics studied. To investigate the impact of maternal antibiotic treatment on the milk metabolome, we profiled the samples via untargeted LC-MS/MS metabolomics. For the metabolomics analyses, all features annotated as antibiotics were removed before conducting comparative analyses (ABX vs. No ABX/Baseline) to limit bias from the antibiotic features themselves. Unsupervised principal component analysis (PCA) revealed separation between groups (PERMANOVA; *pseudo-F*=1.4, *p*=0.002; **Fig. 1e**). Additional covariates influencing the milk metabolome included household income (*F*=1.4) and BMI (*F*=1.3) (all *p* <0.05). To identify metabolic features associated with ABX treatment, we constructed two separate supervised partial least square discriminant analysis (PLS-DA) models comparing ABX vs. Baseline (balanced error rate [BER]=0.41) and ABX vs. No ABX (BER=0.48). Across both models, we identified 633 metabolic features with variable importance projection score (VIP)>1 and concordant directionality; 280 commonly enriched and 353 commonly depleted in response to ABX (**SI Table 1**). To investigate if the differentially abundant features have been previously identified in microbial monocultures, we searched their MS/MS spectra via microbeMASST^33^. Twenty-seven percent had a match, suggesting that these molecules may be produced by bacteria. The natural log ratio of the top 100 features enriched or depleted with ABX significantly separated the ABX group from both Baseline and No ABX (**Fig. 1f**, linear mixed effect model (LME); *p*=1.9e-06 and *p*=0.0003). Acylcarnitines, key components of human milk with essential roles in fatty acid oxidation and energy metabolism^34^, were among the discriminant features associated with ABX exposure, including features with MS/MS matches to isovalerylcarnitine, carnitine-C5:0;O and carnitine-C5:1 (**Extended Data Fig. 1a**). To further contextualize these molecules, we also searched them against tissueMASST^35^ and microbiomeMASST^36^. In addition to matching to microbial monocultures, their MS/MS spectra were also detected more frequently in human blood and urine samples of subjects with chronic disease, such as diabetes mellitus or osteoarthritis, compared to healthy (**Fig. 1g, Extended Data Fig. 1b**). Data from microbiomeMASST using *in vitro* gut synthetic microbial communities (SynCom) further showed that levels of carnitine-C5:0;O decreased during culturing compared to the media control (**Extended Data Fig. 1c**), suggesting deconjugation or dehydroxylation by bacteria. The higher levels of carnitine-C5:0;O in the ABX group could therefore reflect reduced microbial utilization during antibiotic treatment. In contrast, putative linoleic acid and microbially-derived linoleic acid-derived oxylipins (MS/MS matches to 13-Oxo-9Z,11E-octadecadienoic acid, 9-oxo-10E,12Z-octadecadienoic acid, and (9Z,12Z)-15,16-dihydroxyoctadeca-9,12-dienoic acid, but such reference library matches generally do not provide regiochemical or stereochemical elucidation), were depleted in response to ABX. These compounds are important lipid mediators involved in immune modulation^37^. Taken together, these observations suggest that maternal antibiotic treatment may be reflected in altered milk composition, specifically in molecules associated with host-microbial metabolism.

### The infant gut microbiome is taxonomically distinct and enriched in AMR genes

Fecal metagenomics data (*n*=572) were quality filtered (see **Methods**) and downstream analysis was conducted on 554 samples and 915 operational genomic units (OGUs), with a median sequencing depth per sample of 9,328,792 reads. Ordination via phylogenetic-robust principal component analysis (Phylo-RPCA) highlighted a clear separation between the maternal and infant microbiome (**Fig. 2a**, PERMANOVA; *pseudo-F*=709.76, *p*<0.001). Extraction of top taxa differentiating the two groups via PLS-DA (BER=0.05) showed that infants were mainly enriched in *Enterobacteriaceae*, such as *Klebsiella* and *Escherichia*, and *Bifidobacteriaceae*, while mothers presented more *Lachnospiraceae*, *Ruminococcaceae*, and *Oscillospiraceae* (**SI Table 2**). Alpha diversity, measured via Faith’s PD, was higher in mothers compared to infants (**Fig. 2b**, LME; *p*<2.2e-16). Additionally, AMR gene analysis, conducted via the Resistance Gene Identifier (RGI) pipeline against the Comprehensive Antibiotic Resistance Database (CARD)^38^, identified at Baseline a higher proportion of AMR mapping reads in infants compared to mothers (**Fig. 2c**, LME; *p*=3.07e-06) and a direct positive correlation between the two (LM; β = 1.056 and *p*=1.21e-04). Infants are known to harbor more AMR genes compared to adults^39^ that can be vertically or horizontally transmitted from mothers to infants^40^.

**Fig. 2:**
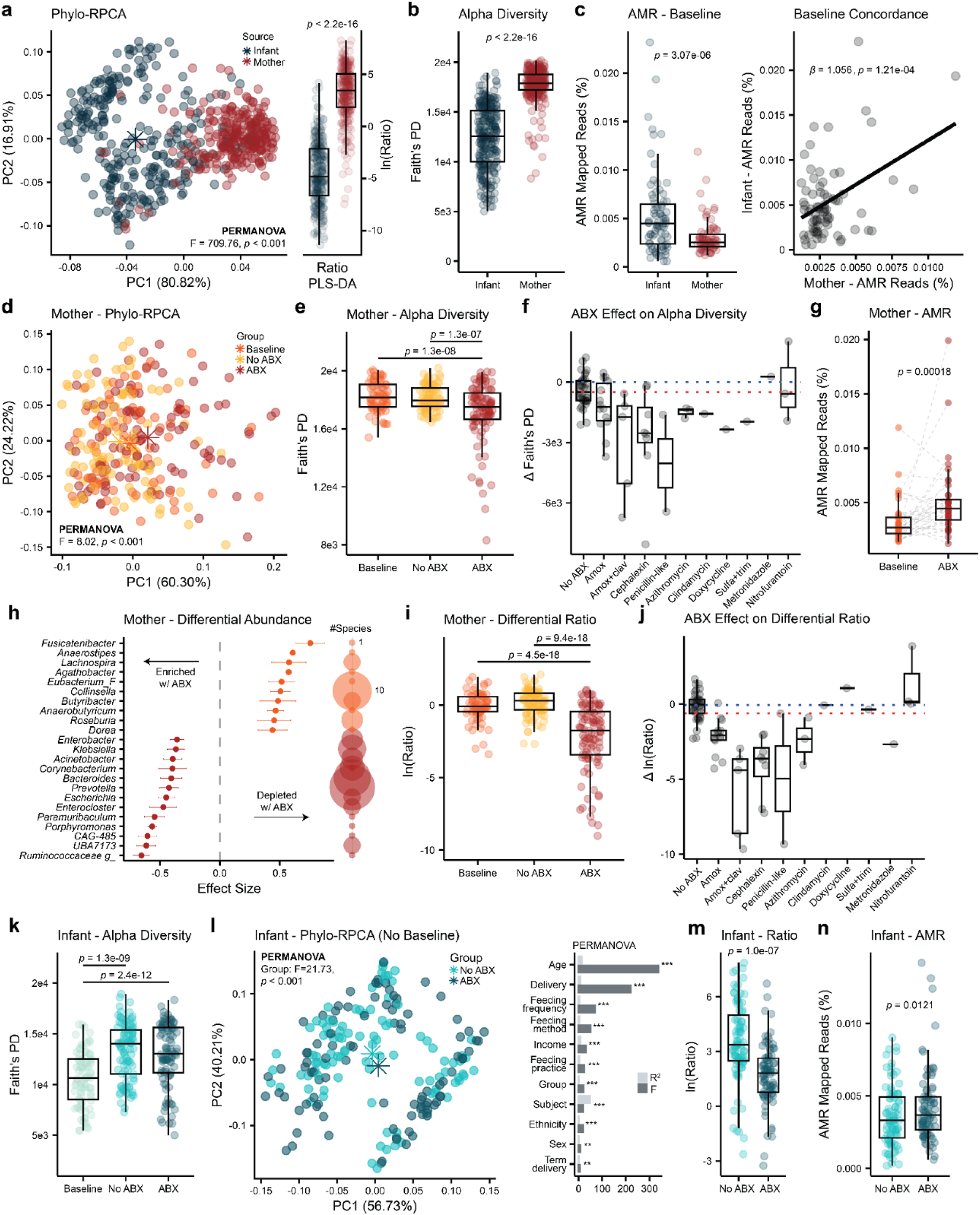
Maternal antibiotic exposure is associated with alterations in both mother and infant gut microbiomes. **a,** Phylo-RPCA of fecal samples separated infant and maternal gut microbial profiles (PERMANOVA; *p*<0.001). PLS-DA showed strong discriminative performance (BER=0.05) and natural log ratio of extracted discriminatory features enriched, at denominator, or depleted, at numerator, in infants recapitulated separation (LME; *p*<2.2e-16). **b,** Mothers presented higher alpha diversity (Faith’s PD) compared to the infants (LME; *p*<2.2e-16). **c,** At baseline infants had a higher relative proportion of AMR mapping reads compared to mothers (LME; *p*=3.07e-06). The proportions of AMR reads between mother and infants were correlated (LM; β=1.056, *p*=1.21e-04). **d,** Phylo-RPCA of maternal samples showed separation between groups (PERMANOVA; *p*<0.001). **e,** Mother treated with ABX presented a lower alpha diversity (Faith’s PD) compared to both Baseline and No ABX groups (LME; *p*=1.3e-08 and *p*=1.3e-07). **f,** Delta between alpha diversity observed at baseline and the subsequent lowest observed value per individual. **g,** Antibiotic treatment increased the proportion of AMR mapping reads from baseline (Wilcoxon Signed-Rank Test, *p*=0.00018). **h,** Differential abundance analysis via ALDEx2 was conducted at Species level by intersecting results from two separate models (ABX vs Baseline and ABX vs No ABX). Only one sample per subject per group was included (see **Methods** for details). Balloon plot was used to visualize the number of differential species present in each genus. **i,** Differential species were used to generate discriminant natural log ratios on all samples, with ABX enriched species at the denominator and depleted ones at the numerator (LME; *p*=4.5e-18 and *p*=9.4e-18). **j,** Delta between ratios observed at baseline and subsequent lowest observed value highlighted a stronger effect of amoxicillin (with or without clavulanate), cephalexin, a penicillin-type drug, and azithromycin. **k,** Infant samples collected at baseline presented a lower alpha diversity (LME; *p*=1.3e-09 and *p*=2.4e-12). **l,** Phylo-RPCA of infant samples without baseline. PERMANOVA identified a separation based on indirect antibiotic exposure (p<0.001), with age, delivery mode, and breastfeeding practice, -method, and -frequency also having an effect. **m,** PLS-DA model (BER=0.29) constructed only on infants indirectly exposed to amoxicillin, cephalexin, a penicillin-type drug, and azithromycin and their matching controls. Differential features were extracted to generate discriminant natural log ratios (LME; *p*=1.0e-07). **n,** Infants indirectly exposed to ABX presented a higher proportion of AMR mapping reads compared to the unexposed counterpart (LME; *p*=0.0121).

### Direct antibiotic exposure is associated with changes in the maternal gut microbiome

Maternal antibiotic treatment was associated with alterations in the overall maternal microbial community structure (**Fig. 2d**, PERMANOVA; *F*=7.99, *p*<0.001) and reduced alpha diversity compared to both Baseline and No ABX (**Fig. 2e**, LME; *p*=1.3e-08 and *p*=1.3e-07). Delta between Baseline and the lowest Faith’s PD value observed in the following timepoints identified amoxicillin (with or without clavulanate), cephalexin, and penicillin-like drug as the antibiotics having the largest impact on alpha diversity (**Fig. 2f**). Additionally, ABX treatment resulted in increased AMR mapping reads compared to Baseline (**Fig. 2g**; Wilcoxon Signed-Rank Test; *p*=0.00018). Multivariate PLS-DA separated the ABX group from both Baseline and No ABX (BER=0.25 and BER=0.20, respectively). A total of 203 microbial species with VIP>1 were commonly enriched (*n*=77) or depleted (*n*=126) in response to ABX compared to both Baseline and No ABX (**SI Table 3**). Additionally, differential abundance analysis via ALDEx2 also identified common enrichment (*n*=106) or depletion (*n*=81) of several bacterial species in response to ABX compared to both Baseline and No ABX (**SI Table 4**), partially recapitulating PLS-DA observations. Importantly, *Bacteroidaceae*, including *Prevotella*, *Bacteroides*, and *Phocaeicola*, *Enterobacteriaceae*, such as *Klebsiella*, *Enterobacter*, and *Escherichia*, *Muribaculaceae*, like *UBA7173*, *CAG-485,* and *Paramuribaculum*, *Porphyromonas*, *Enterocloster*, *Corynebacterium*, and *Acinetobacter* were enriched in response to ABX. On the other hand, *Lachnospiraceae*, such as *Fusicatenibacter*, *Anaerostipes*, *Lachnospira*, *Agathobacter*, *Eubacterium*, *Butyribacter*, *Anaerobutyricum*, *Roseburia*, and *Dorea* species, and *Collinsella* were depleted in response to ABX (**Fig. 2h**). This is in line with previous observations, as the use of β-lactam antibiotics has been associated with reduction in *Lachnospiraceae* and enrichment in *Enterobacteriaceae* and *Bacteroidaceae*^41^. The log ratio of species depleted, at numerator, and enriched, at denominator, with ABX separated the ABX group from both Baseline and No ABX (**Fig. 2i**, LME; *p*<9.4e-18). The change (Δ) between Baseline and the lowest value observed in the following timepoints identified amoxicillin (with or without clavulanate), cephalexin, a penicillin-type drug, and azithromycin as the antibiotics with the largest effect on the maternal microbiome (**Fig. 2j**). The mother receiving metronidazole did not breastfeed her infant during treatment.

### Indirect antibiotic exposure is associated with alterations in the infant gut microbiome

Infant fecal samples showed a lower alpha diversity at Baseline, but no differences between ABX and No ABX (**Fig. 2k**, LME; *p*<1.3e-09). This is likely because infant alpha diversity, unlike maternal alpha diversity, positively correlated with age (LME; β=538.84, *p*<2e-16) and Baseline samples were collected ∼4 months before treatment. Ordination via Phylo-RPCA identified a significant association between indirect ABX exposure and the infant gut microbiome, with or without the inclusion of Baseline samples (**Fig. 2l**, PERMANOVA; *F*=26.46 and *F*=21.73, both *p*<0.001). Nevertheless, covariates with the largest influence on the infant microbiome were age (*F*=337.98) and delivery mode (*F*=219.92), with feeding practices also showing an impact. To further investigate microbial communities associated with indirect ABX exposure, we focused on non-baseline samples collected from infants exposed to either amoxicillin (with or without clavulanate), cephalexin, an unspecified penicillin-type drug, or azithromycin and their matching unexposed controls (*n*=169). This selection was based on the observation that these antibiotics were most strongly associated with alterations in the maternal microbiome. PLS-DA identified a separation between ABX and No ABX (BER=0.29) and 249 microbial species driving it (**SI Table 5**). More specifically, infants indirectly exposed to antibiotics harbored fewer *Prevotella*, *Parabacteroides*, *Alistipes*, *Bacteroides*, *Lactobacillus*, *Alloprevotella*, *Phocaeicola*, and *Bifidobacterium* species and more *Clostridium_A*, *Adlercreutzia*, *Campylobacter*, *Corynebacterium, Blautia*, *Acinetobacter*, and *Enterocloster* species compared to unexposed infants. The depleted genera are known to be altered by β-lactam antibiotics^42^. We observed a reduction of *Prevotella buccalis* and *Prevotella stercorea*, which have been associated with reduced infection risk and improved behavioral development in infants^43,44^, *Parabacteroides johnsonii* and *Parabacteroides goldsteinii*, which have been observed to improve metabolic health, neurocognition, and intestinal barrier integrity in human studies^45^, *Alistipes shahii*, which has been shown to reduce intestinal epithelial damage in animal studies^46^, *Bacteroides salyersiae*, *Bacteroides uniformis,* and *Phocaeicola coprocola*, which can metabolize human milk oligosaccharides (HMOs)^47^ and have been linked to protection of the gut mucosa^48^ and improvement in metabolic health^49^ and neurodevelopment in animal models^50^, *Lactobacillus acidophilus*, *Lactobacillus jensenii*, and *Limosilactobacillus fermentum*, which have been associated with reduced risk of infections^51,52^ and improved immune tolerance^53^, and *Bifidobacterium bifidum*, which can metabolize HMOs and has been shown to positively modulate host metabolic and immune functions^54^. The log ratio of these depleted, at the numerator, and enriched species, at the denominator, also separated ABX from No ABX (**Fig. 2m**, LME; *p*=1.0e-07). Finally, AMR analysis identified a higher proportion of AMR mapping reads in infants indirectly exposed to antibiotics compared to their matched controls (**Fig. 2n**, LME; *p*=0.0121).

### The infant fecal metabolome is characterized by an enrichment of acylcarnitines

In line with the microbiome data, PCA showed a clear separation between infant and maternal fecal metabolic profiles (PERMANOVA; *F*=32, *p*<0.001). PLS-DA (BER=0.002) identified 2,720 differential metabolic features (these could include different adducts or isotopes of the same molecules) with VIP>1 (**SI Table 6**). The natural log ratio of the top 100 enriched and depleted features separated infant and maternal profiles (**Fig. 3a**, LME; *p*<2e-16). Molecular class prediction via CANOPUS^55^ (**Fig. 3b**) identified these features as acylcarnitines (6.2% of all MS/MS spectra of the differential features), bile acids (3.9%), dipeptides (2.9%), aminosugars (2.6%), and amino acids (2.1%). Additionally, 34% (924 features) of them could have putative microbial origin, as they had a match in microbeMASST. By leveraging a recently developed MS/MS library for acylcarnitines^56^ and molecular networking^57^, we identified higher levels of several acylcarnitines, spanning a wide range of chain lengths, unsaturations, and hydroxyl groups, in infants compared to the mothers (**Fig. 3c**). This is consistent with prior studies showing the highest acylcarnitine levels in neonates, followed by a decline during the first year of life to lower levels in adults^58,59^. Additionally, we also observed an enrichment of several candidate bile acids in infants, identified via the detection of the diagnostic fragment ions of the steroid core and unknown modifications^60^.

**Fig. 3:**
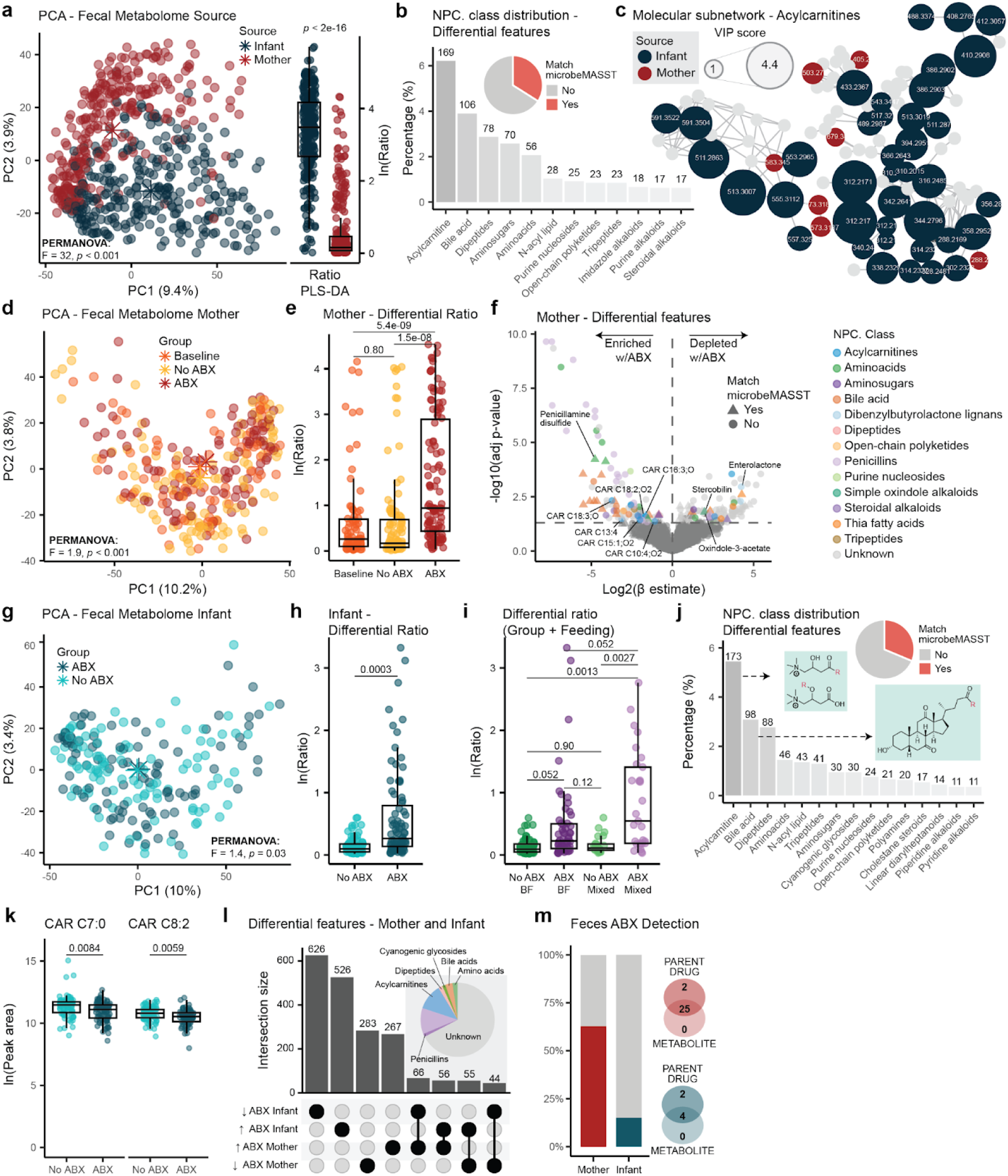
Maternal antibiotic use during lactation is associated with alterations in maternal and infant fecal metabolic profiles. **a,** PCA of fecal samples by source (PERMANOVA; *p*<0.001). Boxplot of the natural log ratio of differential features associated with infant (numerator, top 100) or mother (denominator, top 100) extracted from the PLS-DA model. **b,** Barplot of the distribution of metabolite classes among differential features from the PLS-DA. Features without class prediction and classes with less than 17 features were excluded from the visualization. Pie chart summarizes the proportion of all differential features with a match in microbeMASST. **c,** Submolecular network of differential acylcarnitines. Node size represents VIP scores. **d,** PCA of maternal fecal samples by group (PERMANOVA; *p*<0.001). **e,** Boxplot of the natural log ratio of differential features enriched (numerator, top 100) or depleted (denominator, top 100) with ABX extracted from PLS-DA. **f,** Volcano plot of the differential features identified by PLS-DA. Dots colored by CANOPUS class prediction. Triangle=match in microbeMASST, circle=no match. **g,** PCA of infant fecal samples by group (PERMANOVA; *p*=0.03). **h,** Boxplot of the natural log ratio of differential features enriched (numerator, top 100) or depleted (denominator, top 100) with ABX extracted from PLS-DA. **i,** The log ratio was then stratified by indirect ABX and feeding practices. **j,** Barplot of metabolite class distribution of differential metabolic features from PLS-DA. Features without class prediction and classes with less than 11 features were excluded from the visualization. Pie chart shows the proportion of all differential features with a match in microbeMASST. **k,** Boxplot of selected differential features. **l,** UpSet plot including metabolic features from the PLS-DA models by treatment group for both mother and infant. Only features with mean VIP>1.5 were included in the visualization. The plot summarizes enriched and depleted metabolic features and if they are shared or unique across mothers and infants. **m,** Barplot of detection frequency of antibiotics in fecal samples from the maternal and infant ABX groups. Venn diagrams of the overlap between detections of parent drug only, metabolite only, or both. In **a, e, f, h, i,** and **k** statistical significance was determined using LME with subject as random effect and adjusted for covariates (see **Methods**). All boxplots show the first (lower), median, and third (upper) quartiles, with whiskers 1.5 times the interquartile range.

### Direct antibiotic treatment is associated with different maternal fecal metabolic profiles

Maternal antibiotic treatment was also associated with shifts in maternal fecal metabolic profiles (**Fig. 3d**, PERMANOVA; *F*=1.9, *p*<0.001). PLS-DA models separated ABX from both Baseline (BER=0.40) and No ABX (BER=0.34), and identified 1,284 metabolic features commonly enriched (*n*=636) or depleted (*n*=648) in response to antibiotic treatment (VIP>1, **SI Table 7**). Of these, 32% (409 features) had a match in microbeMASST. The natural log ratio of the top 100 features enriched and depleted with ABX significantly separated the ABX from both Baseline and No ABX (**Fig. 3e**, LME; *p*<5.4e-09 and *p*<1.5e-08). Several acylcarnitines were enriched in fecal samples from the ABX group, particularly medium- and long-chain ones with one or two hydroxy groups and/or unsaturations. These included carnitine-C18:2;O2,C18:3;O, C16:3;O, C10:4;O2, C15:1;O2, and C13:4 (**Fig. 3f**). We also observed shifts in the bile acid pool, including an enrichment of taurine conjugated bile acids in the ABX group, suggesting a decrease in the abundance of bacteria encoding bile salt hydrolase/transferase (*bsh*) and thereby lower capacity for deconjugation^61^, in line with previous findings in murine models^62–64^ (**Fig. 3f**, **SI Table 7**). Additionally, penicillamine disulfide, a breakdown product of penicillin compounds^65^, and other compounds predicted by CANOPUS as penicillins, were enriched in the ABX group (**Fig. 3f**), suggesting the presence of antibiotic-derived drug metabolites. Subsequent investigation of these features revealed almost exclusive detection in fecal samples from amoxicillin-treated mothers (**Extended Data Fig 2a)**. Synthesis of reference standards and MS/MS spectral matching for some of them suggested that these were indeed downstream metabolites of amoxicillin (**Extended Data Fig 2b**). Acetaminophen was also enriched in the ABX group, consistent with symptomatic self-treatment during infection, along with candidate annotations, derived from MS/MS matching, of *N*-acetyl cadaverine-C4:0, urobilin, riboflavin (vitamin B2), and the microbial tryptophan-derived metabolite indole-3-acetic acid (IAA). In contrast, several microbially-derived compounds were depleted during antibiotic treatment, including oxindole-3-acetate, another tryptophan-derived metabolite^66^, the plant lignan enterolactone^67^, and stercobilin, a bilirubin derived metabolite^68^, suggesting shifts in tryptophan and bilirubin metabolism with antibiotic exposure. We also observed a depletion of valine-C3:0, acetylated amino acids, and dipeptides, including putative acetyl-glutamate, acetyl-methionine, acetyl-arginine, acetyl-phe, his-ile, tyr-leu, phe-val, and lac-phe, possibly indicating a broader loss of proteolysis-derived metabolites.

### Indirect antibiotic exposure is associated with different infant fecal metabolic profiles

PCA of the infant fecal metabolome revealed a small, but significant, separation between indirect antibiotic exposure and unexposed matched controls (**Fig. 3g**, PERMANOVA; *F*=1.4, *p*=0.03). Baseline samples, collected on average 4 months prior to treatment, were excluded from downstream analyses due to confounding age effects. Indeed, age was most strongly associated with variation in the infant fecal metabolic profiles (PERMANOVA; *F*=5.3, *p*<0.001), followed by feeding practices (*F*=2.8), type of delivery (*F*=2.3), and delivery mode (*F*=1.8), sex (*F*=1.5), and feeding frequency (*F*=1.4). To identify metabolic features associated with indirect ABX exposure, we restricted analyses to non-baseline samples from infants exposed to either amoxicillin (with or without clavulanate), cephalexin, a penicillin-type drug, or azithromycin and their matching unexposed controls (*n*=170), as for the microbiome analysis. PLS-DA (BER=0.46) identified 3,178 metabolic features enriched (*n*=1,502) or depleted (*n*=1,676) with indirect ABX exposure (**SI Table 8**). The natural log ratio of the top 100 features enriched and depleted with indirect ABX separated ABX from No ABX (**Fig. 3h**, LME; *p*=0.0003). Given that feeding practice was an important covariate, we further stratified the indirect ABX exposure (yes/no) for feeding practice (exclusively breastfed/mixed fed). Notably, we observed that infants receiving a combination of human milk and formula presented a tendency to exacerbated ratios compared to the exclusively breastfed ones (LME; *p*=0.052, **Fig. 3i**). Interestingly, formula feeding is known to differentially influence infant microbiome composition and function^11,69^.

The majority of annotated differential features from the PLS-DA models included acylcarnitines, bile acids, dipeptides, amino acids, and *N*-acyl lipids (**Fig. 3j**). Notably, several short- and medium-chain acylcarnitines, including C7:0, C8:2, C10:1, C14:2;O2, C12:2;O, C6:1, and C8:2;O, were depleted in infants indirectly exposed to ABX (**Fig. 3k, SI Table 8**). Other altered compounds included several known and putative microbial metabolites (31%, 995 features match in microbeMASST). The microbially-derived bile acids, putative Arg-CA and Thr-CA, *N*-acyl lipids, such as histamine-C4:0 and glutamate-C4:0, riboflavin, and IAA were depleted. Indole derivatives, such as IAA, play important roles in maintaining intestinal^70^ and immune homeostasis^71^ by acting as ligands for the aryl hydrocarbon receptor (AhR)^72,73^. In contrast, we observed an enrichment of features that had MS/MS matches to acylcarnitines, such as C12:3 and C15:1;O2, trihydroxylated bile acids, such as Lys-CA and GABA-CA, urobilin, acetyl-arginine, tryptamine-C2:0, arg-C4:0, histamine-C5:0, histamine-C6:0, Tyr-C6:0, Tyr-C7:0, and polyamine derived *N*-acyl lipids, such as cadaverine-C5:0 and cadaverine-C6:0. Several of the *N*-acyl lipids are of microbial origin^74^, and their abundance have been reported to increase rapidly with gut maturation^5^, with cadaverine conjugates having immune modulating effects on CD4+ T cells^74^.

When intersecting features associated with ABX across mothers and infants, we found that the majority of the identified differential features were unique to mothers or infants, suggesting that antibiotic exposure may have distinct effects on adult versus early-life metabolic profiles. A total of 362 metabolic features were shared, of which 166 displayed the same directionality, 96 enriched and 70 depleted (**Fig. 3l**, **SI Table 9**). Compounds commonly enriched in response to ABX included urobilin, CAR-C12:3, CAR-C18:3;O2, CAR-C15:1;O2, and acetaminophen. On the other hand, stercobilin was commonly depleted. Interestingly, several acylcarnitines (such as carnitine-C13:0, C14:1;O, C14:1;O2, C15:2;O, C10:3, C12:1, C9:0), dipeptides (glu-val, val-val, ile-glu, tyr-leu), acetylated amino acids (acetyl-arg), riboflavin, and IAA had opposite directionality between infants and mothers. Finally, we used recently developed MS/MS drug spectral libraries^75,76^ for untargeted metabolomics data to check whether the antibiotics or their downstream metabolites could be detected in the fecal samples. These molecules were detected in 63% of treated mothers and 15% of their infants, with both the parent compound and respective drug metabolites identified in most cases (**Fig. 3m**). Detection of antibiotics in the feces of a fraction of the indirectly exposed infants suggests that antibiotics can be transferred to the infant gut and are probably present at low concentrations around the limit of detection.

### Altered maternal and infant skin metabolomes are associated with maternal antibiotic treatment

Systemic antibiotic treatment during the first year of life nearly doubles the risk of developing atopic dermatitis, the most frequent chronic inflammatory skin disease in both infant and adult populations^77^. For this reason, we assessed if antibiotic exposure was associated with changes in the skin metabolic profiles. PCA showed separation between the maternal and infant skin metabolome (**Fig. 4a**, PERMANOVA; *F*=14.8, *p*<0.001). Via PLS-DA, 1,875 discriminant features were identified (BER=0.007, VIP>1), where 1,129 were enriched and 746 were depleted in infants relative to mothers (**SI Table 10**). The natural log ratio of the top 100 features enriched in mother or infant confirmed significant discrimination (**Fig. 4a**, LME; p<2.2e-16). Briefly, infant skin metabolic profiles presented more compounds annotated as saccharides, while mothers showed more dipeptides. When assessing the association between maternal antibiotic treatment and the maternal skin metabolome, we observed a significant separation via PCA (**Fig. 4b**, PERMANOVA; *F*=1.2, *p*=0.025). Still, ethnicity (*F*=3.2), BMI (*F*=2.4), household income (*F*=2.3), and age (*F*=1.7) were more strongly associated with variation in the maternal skin metabolome (all *p*≤0.006). Additionally, using two PLS-DA models comparing ABX vs. Baseline (BER=0.48) and ABX vs. No ABX (BER=0.43), we extracted 713 metabolic features commonly enriched (*n*=382) or depleted (*n*=331) with ABX (**SI Table 11**). Several of the differential features were dipeptides, either enriched (ile-leu, pro-phe, phe-leu, leu-phe, ile-tyr, arg-ile) or depleted (trp-glu, met-leu). Additionally, carnitine-C5:0 and acetaminophen were enriched with ABX, while phe-C2:0 and the tryptophan derivative glucopyranosyl-tryptophan were depleted. The natural log ratio of features enriched (top 100) and depleted (top 100) with ABX differentiated the ABX group from both Baseline and No ABX (**Fig. 4c**, LME; *p*=0.00013 and *p*=0.0029). There was no statistically significant association between indirect antibiotic exposure and shifts in the infant skin metabolic profiles in a PERMANOVA (**Fig. 4d**, *F*=1.2, *p*=0.09). Age (*F*=2.2), household income (*F*=2.2), delivery mode (*F*=1.6), sex (*F*=1.5), exclusive breastmilk (*F*=1.5), and breastfeeding frequency (*F*=1.8) significantly explained variance in the data (p≤0.007). Nevertheless, PLS-DA comparing the ABX group with the matched controls (Baseline excluded) revealed 1,920 metabolic features enriched (*n*=989) or depleted (*n*=931) in response to indirect ABX exposure (BER=0.47; **SI Table 12**), and the generated natural log ratio of the top 100 features enriched and depleted with ABX separated the two groups (**Fig. 4d**, LME; *p*=2.4e-07). To account for the performance of the skin PLS-DA models and increase confidence in the extraction of features associated with ABX, we intersected the differential features identified in mothers and infants, resulting in 147 metabolic features commonly enriched or depleted across both (**SI Table 13**). The natural log ratio of these shared features significantly separated the maternal ABX group from both Baseline and No ABX (LME; *p*=1.4e-05 and *p*=8.9e-06) and the infant ABX group from No ABX (**Fig. 4e**, LME; *p*=0.0043). Among the ones with a CANOPUS class prediction, we observed dipeptides (5.4%), *N*-acyl ethanolamines (4.1%), *N*-acyl lipids (2.7%), tripeptides (2.7%), amino acids (2.0%), and fatty alcohols (2.0%). Interestingly, 48.3% (71 features) of them found a match in microbeMASST, suggesting important influence of microbial communities (**SI Table 13**). Notably, acetaminophen was enriched in the ABX group in both mothers and infants, while dipeptides such as trp-glu and met-leu were depleted. Finally, we investigated the presence of antibiotics on the skin of treated mothers and their infants. Although antibiotic exposure was associated with alterations in skin metabolic profiles, antibiotics were only detected in 20.9% and 12.5% of maternal and infant skin samples, respectively (**Fig. 4f**).

**Fig. 4:**
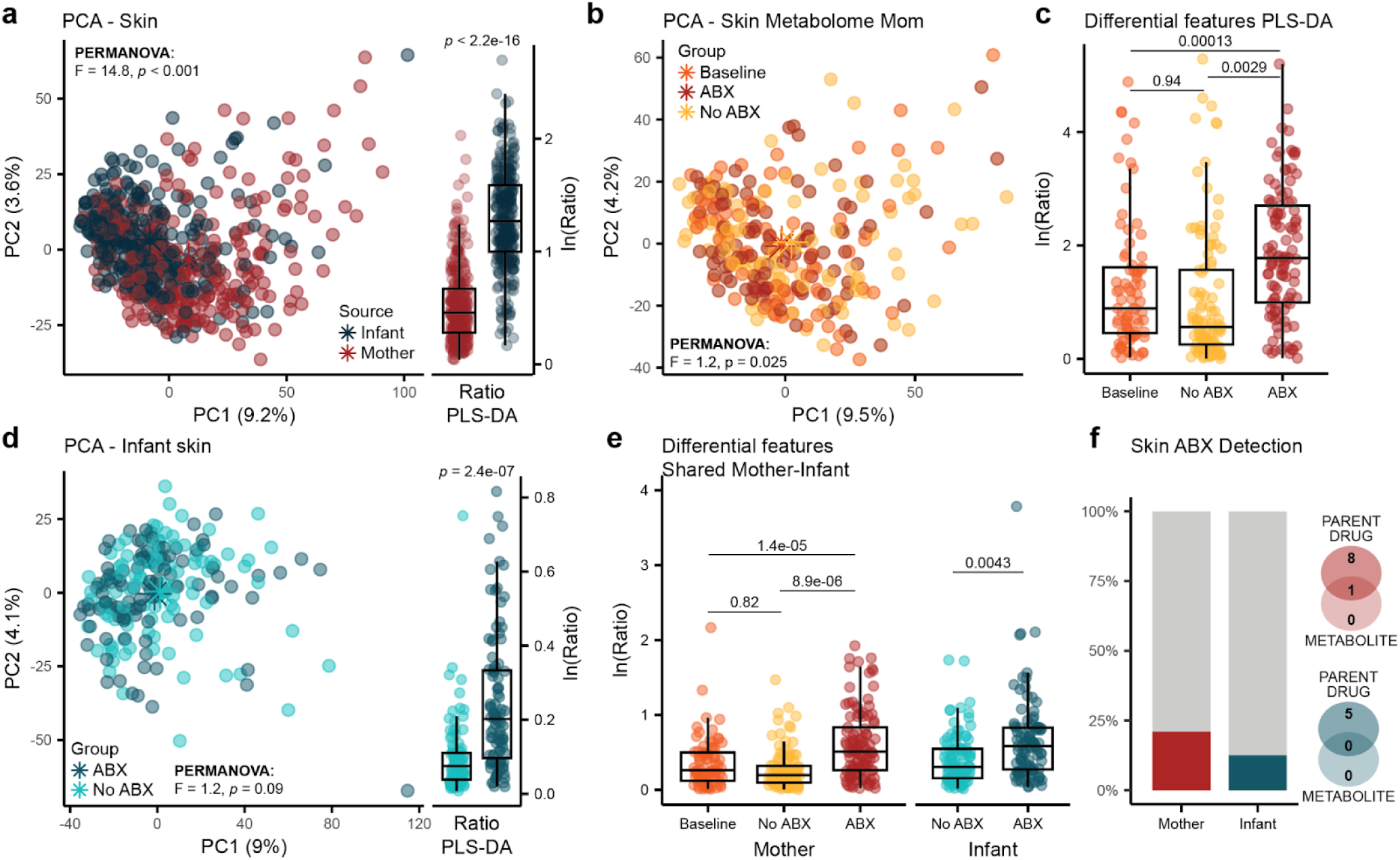
Maternal antibiotic treatment is associated with alterations in skin metabolic profiles. **a,** PCA of skin metabolic profiles by source (PERMANOVA; *p*<0.001), and boxplot of the natural log ratio of differential features enriched in infants (numerator, top 100) or enriched in mothers (denominator, top 100) obtained from PLS-DA (LME; *p*<2.2e-16). **b,** PCA of maternal skin metabolic profiles by group (PERMANOVA; *p*=0.025). **c,** Boxplot of the natural log ratio of differential features enriched (numerator, top 100) or depleted (denominator, top 100) with ABX extracted from the PLS-DA model. **d,** PCA of infant skin metabolic profiles by group (PERMANOVA; *p*=0.09), and boxplot of the natural log ratio of differential features enriched (numerator, top 100) or depleted (denominator, top 100) with ABX from the PLS-DA model. **e,** Natural log ratio of commonly enriched (*n*=61) or depleted (*n*=86) antibiotics-associated features identified by intersecting differential features from the maternal and infant PLS-DA models. **f,** Barplot of antibiotics detection frequency in skin from the maternal and infant ABX groups. Venn diagrams of the overlap between detections of parent drug only, metabolite only, or both. Adjusted p values in **a, c, d,** and **e** were derived from LME with subject as random effect. All boxplots show the lower, median and upper quartiles, with whiskers 1.5 times the interquartile range (IQR).

### Integrative multi-omics analysis and infant anthropometrics

First, we investigated if the ABX-discriminative log ratios generated separately from the microbiome and metabolomics analyses were concordant. We observed a positive correlation in both mothers and infants (**Fig. 5a**, LME; β=0.51, *p*<2e-16 and β=0.16, *p*=0.012). Then, we performed multi-omics integrative analyses via Joint-RPCA on maternal and infant samples separately. Heatmaps showing associations between annotated metabolites and bacterial species of interest were generated for both mothers and infants (**Fig. 5b** and **Fig. 5c**). Importantly, medium- and long-chain carnitines correlated with several *Enterobacteriaceae*, including *Klebsiella*, *Enterobacter*, and *Escherichia*, in maternal fecal samples and were enriched, like these taxa, in response to antibiotics. Previous research supports this observation, as carnitines can promote the growth of *Enterobacteriaceae*^78^ and perinatal administration of ampicillin, another broad-spectrum β-lactam antibiotic, was shown to induce fecal enrichment of carnitines and *Enterobacteriaceae* in a murine model^79^. Interestingly, short-and medium-chain carnitines were depleted with indirect exposure to antibiotics in infants and they were correlated with taxa considered beneficial, such as *Bifidobacterium*, *Bacteroides*, *Prevotella*, and species belonging to the *Muribaculaceae* family. Interestingly, infants indirectly exposed to antibiotics presented higher carnitine C12:3, which was detected at a higher frequency in inflammatory conditions via tissueMASST (**Fig. 5d**). Antibiotic exposure was also associated with alterations in heme catabolism in both mothers and infants, with enrichment of urobilin and a depletion of stercobilin in both. Bilirubin detoxification is catalyzed by the gut microbiome, with bilirubin first being converted to urobilinogen and then to stercobilin for excretion^80^. As Bacillota, formerly Firmicutes, is the main phylum encoding this metabolism, we observed in mothers a strong correlation of stercobilin with *Lachnospiraceae* species, also depleted in response to antibiotics. Finally, glutamate-C4:0 was associated in infants with *Bifidobacterium* species, which are known metabolizers of glutamate^81^.

**Fig 5.**
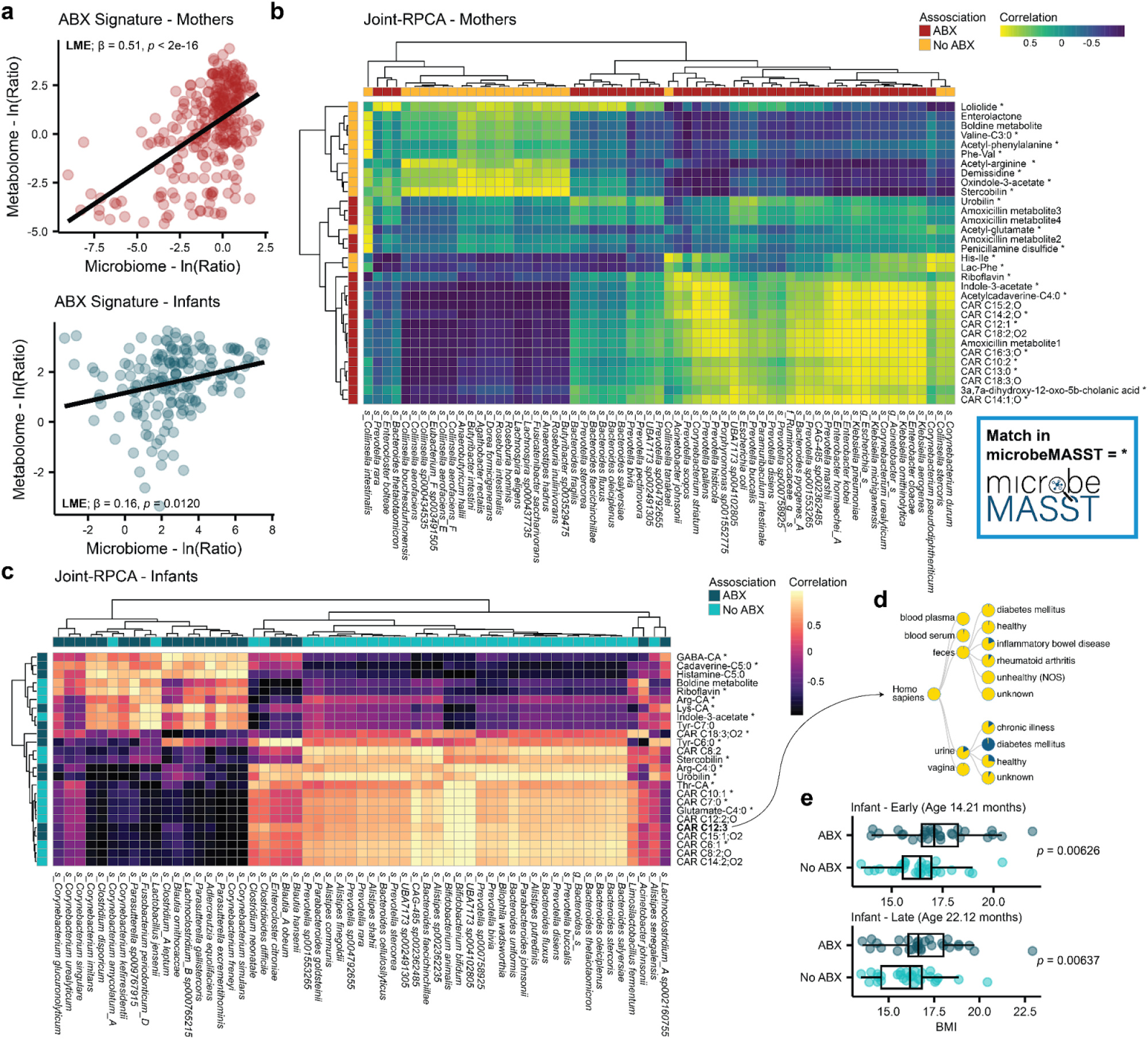
Multi-omics integration of maternal and infant fecal samples. **a,** Correlation between identified ABX-discriminative log ratios in the microbiome and metabolomics analyses for mother and infants (LME; *p*<2e-16 and *p*=0.0120). **b,** and **c,** Heatmaps generated from correlations tables of the Joint-RPCA analysis. Selected microbial and metabolic features of interest are visualized. Information if the features were associated with antibiotic exposure in the separate single omics analysis is reported. Matches in microbeMASST are reposted as *. **d,** Search output from carnitine C12:3 in tissueMASST. **e,** Boxplots from BMI measurement at 14.2±6.0 months (Early) and 22.1±5.6 months (Late) of age in a subset of infants (*n*=61, LM; p<0.0063).

Anthropometric data were collected for a subset of infants (*n*=61). Importantly, infants that were indirectly exposed to antibiotics presented higher BMI compared to their matched controls, after adjusting for age, delivery mode, sex, term delivery, and feeding practices (LME, *p*=0.0187). Additionally, we stratified BMI analysis based on earliest recorded measurement (mean age in months: 14.2±6.0) or latest (mean age: 22.1±5.6) and still observed a separation between the two groups after adjusting for covariates (**Fig. 5e**, LM; *p*<0.0063). Direct early-life exposure to antibiotics has been associated with increased BMI^82^, which can then increase susceptibility to develop metabolic diseases later in life^83^.

## DISCUSSION

Breastfeeding represents the gold standard to support infant development and the gut microbiome^84,85^, but potential disruptions associated with maternal antibiotic treatment during lactation remain largely unexplored. Here, we show that maternal antibiotic treatment is associated with alterations in the maternal gut microbiome and metabolome, that antibiotics are secreted into human milk, and that this indirect exposure may be associated with changes in the gut microbiome composition and function of breastfeeding infants.

Antibiotics were consistently detected in milk samples. Azithromycin and trimethoprim had the highest concentrations, in line with their physicochemical properties facilitating excretion into milk compared to more hydrophilic antibiotics, such as amoxicillin^20^. Timing of milk collection in relation to antibiotic exposure and interindividual differences in pharmacokinetics may also contribute to the observed variations. Although antibiotic use showed weak association with altered milk metabolic profiles, we identified differences in putatively annotated microbially-derived metabolites, such as an enrichment of carnitine-C5:0;O and carnitine-C5:1. While the biological role of these molecules remains unexplored, hydroxyvalerate (C5:0;O) and tiglic acid (C5:1) can derive from microbial metabolism of leucine and isoleucine, two branched-chain amino acids (BCAAs)^86^. Bacteria encoding for these metabolic pathways, such as *Bacteroides* and *Prevotella* genera^87,88^, were enriched in the maternal fecal microbiome in response to treatment. Additionally, carnitines are involved in β-oxidation and represent a fundamental source of energy for the host^34^. As such, when transferred to infants, these molecules can influence the developing metabolic health.

As expected, antibiotic treatment was strongly associated with changes in the maternal fecal microbiome composition and activity. Together with a reduction in alpha diversity, we observed a depletion of abundant commensal taxa that can produce butyrate, such as *Fusicatenibacter saccharivorans*, *Lachnospira eligens*, *Anaerostipes hadrus*, *Agathobacter rectalis*, and *Anaerobutyricum hallii*^89,90^. Additionally, antibiotic exposure significantly increased the proportion of AMR mapping reads in the maternal microbiome, which could be deleterious both to the mother and to the infants, in case of vertical or horizontal transfer. Maternal antibiotic treatment resulted in an enrichment of *Enterobacteriaceae*, such as *Klebsiella*, *Enterobacter*, and *Escherichia*, and several medium- to long chain acylcarnitines. Multi-omics integration via Joint-RPCA showed a strong correlation between *Klebsiella* and *Enterobacter* species and these carnitines; a relationship repeatedly observed in human and animal studies^78,79^.

Furthermore, we identified a significant association between indirect antibiotic exposure and changes in the infant fecal microbiome and metabolic profiles. Although age, mode of delivery, and feeding practices were more strongly associated with variation in the composition of the fecal microbial communities, infants indirectly exposed to antibiotics presented lower levels of *Prevotella, Parabacteroides, Alistipes, Bacteroides, Lactobacillus, Phocaeicola, and Bifidobacterium*. As these taxa are typically depleted during treatment with β-lactam antibiotics, exposure via maternal milk might represent the cause of their disruption. Importantly, the higher proportion of AMR gene reads in infants indirectly exposed to antibiotics suggests that even low-dose exposure may exert selective pressure on the infant resistome, potentially increasing the risk of treatment failure should these infants require future antibiotic treatment. This is in line with previous reports showing a higher occurrence of β-lactamase coding genes in infants where the mother received intrapartum antibiotic prophylaxis^91^.

Interestingly, formula feeding, known to influence the early-life fecal microbiome^8^, appeared to exacerbate the metabolic alterations associated with ABX exposure, suggesting shared cumulative effects. Furthermore, we observed a depletion of short- and medium-chain acylcarnitines in infants indirectly exposed to antibiotics. This may reflect a lower availability of microbial short-chain fatty acids (SFCA) to be conjugated with carnitine, consistent with the observed depletion of taxa producing them, such as *Bacteroides, Bifidobacterium,* and *Prevotella*. Given that acylcarnitine levels were shown to be more abundant in infants than in adults^58,59^, infants may be more sensitive to altered levels of their microbial precursors. We also observed associations between antibiotic exposure and heme catabolism, characterized by an enrichment of urobilin and a depletion of stercobilin in response to ABX exposure. This suggests a preferential metabolism of the common intermediate urobilinogen to urobilin rather than stercobilin^80^. Although the enzyme responsible for this transformation remains unknown, this might reflect a depletion of taxa performing this conversion. Further, we observed a depletion of indole-3-acetic acid, a metabolite whose dysregulation has been associated with different health conditions, including persistent overweight from childhood into adolescence^92^. Several short-and medium-chain *N*-acyl lipids were also increased in response to ABX. Specifically, cadaverine conjugates, which have been shown to have immune modulating effects and to be associated with cognitive impairment in adults^74^. Additionally, higher levels of *N*-acyl lipids have been observed in infants born via C-section during the first month of life^5^ and in children diagnosed with very early onset inflammatory bowel disease (VEO-IBD)^93^, although these compounds presented different head groups.

Finally, indirect antibiotic exposure was associated with a higher BMI in a subset of infants with available anthropometric information. To support the idea that even low-level antibiotic exposure during the perinatal developmental window can have a long-term impact on infant health, a previous retrospective study conducted on 200,000 infants showed that intrapartum antibiotic prophylaxis for Group B *Streptococcus* (GBS-IAP) was associated with sustained BMI increase from infancy through childhood^94^. Importantly, higher BMI during childhood has been linked to an increased risk of developing non-communicable diseases, such as type 2 diabetes, later in life. Although maternal infection status *per se* remains a potential confounder, antibiotic-induced alterations could be a more plausible mechanism than underlying inflammation, given the established strong and prolonged effect of antibiotics on the gut microbiome^95^.

In conclusion, we show that maternal antibiotic treatment, often necessary during lactation, is associated with changes in both the maternal and infant microbiome. Nevertheless, further research is required to understand how this could potentially affect long-term health outcomes of both mothers and developing infants. Validation in larger cohorts will provide guidance for antibiotic treatment during lactation as well as infant feeding practices during this period.

### Limitations

Antibiotic treatment was based on clinical indication, with the majority of the mothers receiving amoxicillin (with or without clavulanate) or cephalexin. As such, findings should be interpreted primarily as an effect of β-lactam antibiotics, and may not be directly generalizable to other antibiotic classes. Although the study included age-matched controls and age was added as a covariate in all the models, this variable exerted a substantial influence on both the microbiome and metabolome of the infants. Larger or stricter age-defined studies should be performed to confirm observations. Short-read metagenomic sequences were aligned to reference databases and taxonomic assignments at the species level should be interpreted with caution due to inherent sequence homology among closely related taxa. Future studies should implement long-read sequencing to generate high-quality metagenome assembled genomes (MAGs) and improve taxonomic resolution. Metabolite annotation was based on MS/MS spectral library matching, which represents a Level 2-3 annotation according to the Metabolomics Standard Initiative (MSI)^96^. Anthropometric data, specifically Body Mass Index (BMI), were available only for a subset of infants and larger infant cohorts should be investigated to confirm these findings. Additionally, long-term health repercussions should be evaluated to identify possible development of metabolic diseases.

## Acknowledgements

We thank Gail Ackermann for assistance in depositing metagenomics data in EBI/ENA. This publication includes data generated at the UC San Diego IGM Genomics Center utilizing an Illumina NovaSeq X Plus that was purchased with funding from a National Institutes of Health SIG grant (#S10 OD026929).

## Author contributions

K.E.K. processed untargeted metabolomics samples, conducted untargeted metabolomics analysis, multi-omics analysis, generated figures, and wrote the manuscript.

S.Z. conducted metagenomics analysis, multi-omics analysis, generated figures, and wrote the manuscript.

K.B. and C.C. conducted the clinical study.

J.C., R.T.S., and J.M. performed targeted quantification of antibiotics in milk.

V.C.L, A.P., V.D., S.X., J.A., J.Z., Y.E.A., S.T., A.G., A.G., A.L., and B.H. processed untargeted metabolomics samples or contributed to untargeted metabolomics analyses.

R.K., C.C., P.C.D., and S.M.T. provided supervision, designed the study, and/or secured funding.

All authors reviewed and approved the manuscript.

## METHODS

### Study Design, Recruitment, and Ethics Statement

Breastfeeding women with a child less than 6 months of age who resided in Southern California and never took antibiotics between the birth of their child and study enrollment were recruited and enrolled into Mommy’s Milk, the Human Milk Research Biorepository (HMB) at the University of California, San Diego between 2022 and 2024. The structure and design of the HMB have been described elsewhere^31^. Women who volunteered for the study were recruited into HMB through a variety of sources, including self-referral, healthcare providers, the MotherToBaby counseling service, research studies, direct-to-consumer advertisement, and social media. Participants provided written informed consent for future research uses of milk samples and associated data. The HMB protocol was approved by the institutional review board at the University of California San Diego, (IRB #130658) and a National Institutes of Health (NIH) Certificate of Confidentiality was obtained.

#### Baseline Samples and Associated Data

Following consent, study participants were mailed sample collection materials and sample collection instructions for their baseline samples which included breast milk, maternal and infant feces, and maternal and infant skin swabs. Within 24 hours of baseline sample collection, breastfeeding mothers completed their baseline interview providing demographics, maternal and child health history, child feeding patterns, child adverse reactions and details regarding exposures to medications, alcohol, tobacco, and other recreational substances for the 14 days prior to milk sample collection. Following baseline sample collection, participants received weekly text messages querying them on continued breastfeeding and any new antibiotic prescriptions. If a participant responded yes to both continued breastfeeding and a new antibiotic prescription, study staff contacted them within 24 hours to enroll into the antibiotic exposed sub-cohort.

#### Antibiotic Exposed and Matched Controls Sub-Cohort

Women who confirmed a systemic antibiotic were enrolled into the antibiotic exposed sub-cohort and were sent additional sample collection materials. Participants were asked to collect up to three sample sets (breast milk, maternal and infant feces and skin swabs) during the course of their antibiotic treatment. Ideal samples were collected at the beginnings of treatment (Day 1-2), middle (Day 3-5), and end (Day 7-10) depending on the specific antibiotic prescribed. Simultaneously, a participant from the baseline cohort was matched to the new antibiotic exposed participant on infant sex and age and contacted for enrollment into the control sub-cohort. The matched controls were asked to collect three sample sets at the same timepoints as their matched counterpart. Within 24 hours of their last sample collection, breastfeeding mothers completed another interview providing updates to their medical history and their child’s health history, child feeding patterns, child adverse reactions and details regarding exposures to medications, alcohol, tobacco, and other recreational substances for the 14 days prior to sample collection.

### Sample Collection

#### Breast Milk

Participants were instructed to pump and collect 50 mL of milk up to a full expression as close to the time of the scheduled interview as possible and up to 24 hours prior. Any quantity of milk ≥1 mL was accepted. Participants used a personal electric breast pump and collection containers during milk collection and then transferred the expressed milk to a study provided sterile milk collection bag. Samples were stored in the home refrigerator until time of shipment. A courier service transported samples overnight on ice to the Mommy’s Milk lab where samples were cataloged and stored at -80 ℃ until analysis.

#### Maternal and Infant Feces

Participants were instructed to rub a provided cotton swab against soiled toilet paper to collect their maternal fecal sample. For the infant fecal sample, participants were instructed to rub a provided cotton swab against the soiled diaper. Participants were asked to saturate at least half of the cotton tip with feces. Samples were then stored in the home refrigerator until time of shipment. A courier service transported samples overnight on ice to the Mommy’s Milk lab where samples were cataloged and stored at -80 ℃ until sample analysis.

#### Maternal and Infant Skin Swabs

Participants were instructed to remove a pre-moistened (50% ethanol) swab from the tube and firmly rub both sides of the cotton swab on the back of the neck in a circular pattern for 30 sec with moderate pressure. The swab was then placed into an empty tube and stored in the home refrigerator until time of shipment. A courier service transported samples overnight on ice to the Mommy’s Milk lab where samples were cataloged and stored at -80 ℃ until sample analysis.

### Antibiotic Quantification in Human Milk

Antibiotics in human milk were quantified by liquid chromatography mass spectrometry (LC-MS). LC separation was performed using a Waters HSS T3 column (1.8 μm, 2.1 mm x 50 mm, SKU: 186003538) on a Waters ACQUITY UPLC I-Class FTN PLUS System. Mobile phase A was 1% formic acid and 99% 18MΩ cm H_2_O. Mobile phase B was 0.1% formic acid and 99.9% methanol. Multiple Reaction Monitoring-Mass Spectrometry (MRM-MS) was performed on a Waters Xevo TQ-XS Triple Quadrupole mass spectrometer and the acquired data was analyzed using TargetLynx. Azithromycin, Sulfamethoxazole, Doxycyclin, Cephalexin, Nitrofurantoin, Trimethoprim, Clindamycin, and Amoxicillin were purchased together with their stable-isotope labelled analogs (Azythromycin-d_3_, Sulfamethoxazole-d_4_, Trimethoprim-d_3_, Doxycyclin-d_3_, Cephalexin-d_5_, Nitrofurantoin-^13^C_3_, Clindamycin-d_3_, and Amoxicillin-d_4_). External calibration curves were generated via isotope dilution using matrix-matched calibrators for each antibiotic. The external calibration curves were subsequently used to quantify antibiotics in breast milk samples.

### Metagenomics Sequencing and Processing

Fecal samples were transferred to Matrix Tubes^®^ (ThermoFisher, USA) for gDNA extraction and sequencing^97^, as previously described^5^. Briefly, this high-throughput protocol uses a MagMAX Microbiome Ultra Nucleic Acid Isolation Kit (ThermoFisher Scientific, USA), a Quant-iT PicoGreen dsDNA Assay Kit (Invitrogen, USA), and a miniaturized adaptation of the KAPA HyperPlus Library Kit (Roche, Switzerland). Samples were equal volume pooled, PCR cleaned (Qiagen, Germany), and size selected from 300 to 700 bp. Expected library sizes were confirmed via an Agilent 4000 Tapestation (Agilent, USA). An iSeq100 (Illumina, USA) was used to sequence the equal volume pool. The obtained read counts per sample were used to optimize pooling efficiency and obtain more even per sample read counts. Sequencing was performed on a NovaSeq X Plus (Illumina, USA) at the UCSD Institute for Genomic Medicine (IGM) using 25B flow cells and 2 × 150 bp chemistry. Raw sequence reads were demultiplexed to per sample FASTQ and quality filtered. After adapter trimming via fastp^98^, human reads were filtered out using a recently developed host filtration pipeline^99^. Briefly, reads mapping to reference human genomes (GRCh38.p14 + Phi X 174 + T2T-CHM13v2.0) via minimap2^100^ were removed. Then, an additional filtering step using a pangenome index generated from 94 reference assemblies from the Human Pangenome Reference Consortium^101^ using Movi^102^ was performed. Cleaned FASTQ files were uploaded into Qiita^103^ (<u>Study ID #15626</u>). Data were processed using Wolkta^104^, via the default workflow and parameters. In short, qp-woltka 2024.09 used bowtie2^105^ (v 2.5.4) to align reads against the Web of Life (WoL2)^106^ reference database and generate operational genomic units (OGUs). These were then filtered against Greengenes2 (gg/2024.09) to generate the final BIOM^107^ table. After extracting the coverage.tgz files from Qiita, micov^108^ was used to apply a coverage dispersion filter and remove low confidence OGUs. OGUs with less than 5% genome coverage were removed, together with OGUs detected in less than 5% of the total samples. Finally, samples with less than 1,000,000 cumulative reads were discarded (*n*=18). Downstream analysis was then conducted on 554 samples and 915 OGUs. Median sequencing depth was 9,161,128 reads. Sequencing depth was significantly higher (Mann-Whitney U Test, *p*<2.2e-16) in infants (median=11,043,556) compared to mothers (median=7,573,525).

### Metagenomics Data Analysis

Downstream analyses were conducted in R 4.5.2 (R Foundation for Statistical Computing, Vienna, Austria), QIIME2^109^ (q2cli v 2025.4.0), and Python 3.10.14 (Python Software Foundation). Greengenes2 was used for taxonomy and phylogeny. Alpha diversity metrics (Shannon and Faith’s PD) were calculated using QIIME2 after a rarefaction step (sampling depth: 1,000,000). The two metrics were strongly correlated (LM; p<2.2e-16). Alpha diversity analysis detected extreme outliers between maternal samples (*n*=3) at Baseline and in No ABX, which were removed from subsequent analysis. Group differences in univariate analyses were evaluated via linear mixed effect (LME) models, to take into account the repeated measures, using ‘lmerTest’ (v 3.2-0)^110^. The package ‘emmeans’ (v 2.0.1) was used to compute estimated marginal means from LME models and pairwise *p* values using Tukey’s HSD for adjustment. For maternal alpha diversity analysis (*n*=272) the following LME model was used: faith_pd ∼ group + age_at_collection + BMI + ethnicity + household_income + (1|subject_id). Additionally, to capture the delta in alpha diversity between baseline and treatment, repeated measures per subject during ABX exposure (or matching control) were filtered to retain the lowest Faith’s PD value observed across samples. These were used to calculate the delta from baseline. In both ABX and No ABX, a significant decrease in alpha diversity from baseline was observed (Wilcoxon Signed-Rank Test, p=2.2e-06 and p=0.0036) but the magnitude of this change was greater in the ABX group (Cliff’s δ [95% CI]: 0.63[0.40, 0.78]) compared to the No ABX group (Cliff’s δ [95% CI]: 0.30[0.03, 0.53]), calculated via ‘effsizè (v 0.8.1). Phylogenetic robust principal component analysis (Phylo-RPCA)^111,112^, a compositionality and phylogeny aware beta diversity metric, was also performed via ‘gemellì in QIIME2, without any rarefaction. Phylo-RPCA was calculated on all samples, mother samples only, and infant samples only (with or without the inclusion of baseline samples). All outputs were exported in R for downstream analyses and visualization. PERMANOVA and PERMDISP2 on the Phylo-RPCA distances matrices were conducted using the functions adonis2 and betadisper from ‘vegan’ (v 2.7-2), respectively. The function decostand was used to perform robust center log ration (RCLR) transformation (without matrix completion; impute=FALSE) on the OGUs tables. PERMANOVA on maternal beta diversity was performed using the following formula: distance_matrix ∼ group + age_at_collection + BMI + ethnicity + household_income + subject_id. Features with near zero variance were removed using ‘caret’^113^ (v 7.0-1) before dimensionality reduction. Principal component analysis (PCA) and partial least square discriminant analysis (PLS-DA) were performed on the caret-cleaned RCLR transformed data via ‘mixOmics’ (v 6.34.0)^114^. PLS-DA models performances were evaluated using 10-fold cross-validation and 999 repeats. Models with balanced error rate (BER) < 0.5 were considered discriminant. Features with variable importance projection (VIP) score > 1 were considered relevant for the discrimination. OGUs tables were manipulated using ‘phyloseq’ (v 1.54.0)^115^ and agglomerated at Species level before differential abundance analysis. ALDEx2^116^ (v 1.42.0) was used to calculate effect sizes and evaluate significant differences between groups. For maternal differential abundance analysis, only one sample per subject, the one with the lowest alpha diversity, was retained to avoid the use of repeated measures. Only microbial species commonly enriched or depleted following ABX exposure, after intersecting results from ABX vs Baseline and ABX vs No ABX models, were considered significant and used to generate the final log ratios. Features enriched in response to antibiotics were summed at the numerator, while the depleted ones were summed at the denominator. Log ratio was calculated on all samples and LME was then used to investigate group differences using this formula: log_ratio ∼ group + age_at_collection + BMI + ethnicity + household_income + (1|subject_id). Effect size between models (ABX vs Baseline and ABX vs No ABX) and of significant species within each genus were collapsed to the mean and standard deviations were calculated. The number of significant species within each genus of interest was calculated and visualized using a balloon plot. For infant data (*n*=273), age represented the strongest source of variation in both alpha and beta diversity analyses. PERMANOVA was conducted using the following formula: distance_matrix ∼ group + age_at_collection + delivery_mode + sex + ethnicity + type_delivery + household_income + exclusive_breastmilk + breastfeeding_method + breastfeeding_frequency + subject_id. As baseline samples were collected on average 4 months before treatment, they were not used as reference points, and downstream investigation was only focused on differences between samples collected during treatment, ABX and No ABX group (*n*=194). Additionally, since the strongest antibiotic effects in mothers were observed for amoxicillin (with or without clavulanate), cephalexin, penicillin-type drug, and azithromycin in mothers, only infants indirectly exposed to these antibiotics and their matching controls were investigated for differential abundance analysis (*n*=169). Features used to generate the final infant differential log ratios were extracted from a PLS-DA model. LME was then used to detect antibiotic effect via the following formula: log_ratio ∼ group + age_at_collection + delivery_mode + sex + exclusive_breastmilk + breastfeeding_frequency + type_delivery + (1|subject_id). Data were manipulated using ‘tidyversè (v 2.0.0) and visualized via ‘ggpubr’ (v 0.6.2).

### Antimicrobial Resistance (AMR) Gene Analysis

Per sample FASTQ files were downloaded from Qiita to perform AMR profiling on the acquired metagenomics data. Files were already cleaned from human reads as described in the previous paragraph. A docker image of the Resistance Gene Identifier (RGI) application (v 6.0.5) was locally deployed on a Linux workstation and both the Comprehensive Antibiotic Resistance Database (CARD) (v 4.0.1) and WildCARD (CARD’s Resistomes & Variants and Prevalence Data) were used as reference databases^38^. The default reads aligner KMA^117^ was used, as recommended. The extracted per sample output (*gene_mapping_data.txt) was then concatenated in R and only genes with an average percent coverage > 30% and present in 5 or more samples were retained for downstream analysis. Group differences were investigated with the same model framework described for the taxonomic analysis, LME when repeated measures were present and Wilcoxon Signed-Rank Test for paired data.

### Untargeted Metabolomics Sample Preparation

A total of 573 fecal, 292 human milk, and 590 skin samples were prepared for untargeted metabolomics analyses. Feces were extracted using 95% ethanol (EtOH) via the Matrix tube method as described previously^97,118^. In brief, samples were extracted in 400 µL of 95% EtOH, shaked at 1,200 rpm for 2 min in a SpexMiniG plate shaker (SPEX SamplePrep part #1600, NJ, USA), centrifuged for 5 min at 2,700 g, and then the supernatant (200 µL) was collected and stored at -80 °C for later acquisition. Before instrumental analysis, samples were reconstituted in 200 µL 50:50 methanol (MeOH):water. Milk was extracted using 75:25 MeOH:methyl tert-butyl ether (MTBE) as described previously^119^. In brief, 50 µL of milk was added to a 96-deep well plate and extracted in 200 µL of cold MeOH/MTBE, sonicated for 5 min, and centrifuged for 15 min at 1800 rpm (4 °C). Subsequently, the supernatant (150 µL) was transferred to a new 96-well plate and analyzed immediately by LC-MS/MS. Swabs used for skin sampling were extracted in 500 µL of 50:50 MeOH:water, sonicated for 5 min, sealed with an aluminium cover, and incubated overnight at 4 °C, as previously described^120^. The following day, swabs were removed from the extracts using pre-cleaned stainless steel tweezers, with rinsing between. The extracts were then concentrated to dryness in a CentriVap and stored at -80 °C until analysis. Prior to LC-MS/MS analysis, samples were reconstituted in 200 µL of 50:50 MeOH:water, sonicated for 5 min, centrifuged for 5 min at 1500 rpm, before the supernatant (150 µL) was collected and used for instrumental analysis. Sulfadimethoxine (1 µM) was used as the internal standard for all sample matrices.

### Untargeted Metabolomics Data Acquisition

Randomized samples (5 μL) were injected into a Vanquish ultra-high-performance liquid chromatography (UHPLC) system coupled to a QExactive quadrupole orbitrap (Thermo Scientific) mass spectrometer. Chromatographic separation was performed on a polar C18 column (Kinetex Polar C18, 100 mm x 2.1 mm, 2.6 µm particle size, 100Å; Phenomenex) with a matching Guard cartridge (2.1 mm). The mobile phase was solvent A (water with 0.1% formic acid) and solvent B (acetonitrile with 0.1% formic acid), and all solvents used were LC-MS grade. The flow rate was set to 0.5 mL/min. For fecal and skin samples, the following gradient was applied: 0-1 min at 5% B, 1-7.5 min to 40% B, 7.5-8.5 min to 99% B, 8.5-9.5 min at 99% B, 9.5-10 min to 5% B, 10-10.5 min at 5 % B, 10.5-10.75 to 99% B, 10.75-11.25 min at 99% B, and 11.25-11.5 to 5% B. For milk samples, the gradient was slightly modified: 0-1 min at 5% B, 1-8.5 min to 99% B, 8.5-9.5 min at 99% B, 9.5-10 min to 5% B, 10-10.5 min at 5% B, 10.5-11 min to 99 % B, 11-11.5 at 99% B, 11.5-12 min to 5% B. Tandem mass spectrometry (MS/MS) data was acquired in data-dependent acquisition (DDA) mode with positive electrospray ionization (ESI+). Source parameters applied: sheath gas flow 53 L/min, aux gas flow rate 14 L/min, sweep gas flow 3 L/min, spray voltage 3.5 kV, inlet capillary 269 °C, aux gas heater 438 °C, and 50 V S-lens level. The MS scan range was set to 150-1500 *m/z* for feces and milk samples, and 100-1000 *m/z* for skin samples, with a resolution of 35,000. Automatic gain control (AGC) target was set to 1e6 with a maximum ion injection time of 100 ms. MS/MS were collected with a resolution of 17,500 and AGC target of 5E5 with a maximum ion injection time of 150 ms. The top five most abundant ions per cycle were fragmented, with isolation window set to 1.0 *m/z*, isolation offset to 0 *m/z*, and stepped collision energies (25, 40, 60 eV). 5 s dynamic exclusion window was applied.

### Untargeted Metabolomics Data Processing

The acquired .raw files were converted into .mzML format using ProteoWizard MSConvert^121^. Data files can be found in <u>GNPS/MassIVE</u> under the accession code <u>MSV000099924</u> (feces), <u>MSV000100756</u> (skin), and <u>MSV000100755</u> (human milk). Feature detection and extraction was performed using MZmine^122^ version 4.7.8, with custom settings for each biofluid. For mass detection, the factor of lowest signal was used with noise factor 5 for MS1 and 2.5 for MS2 for all sample matrices. <u>Feces:</u> parameters in the chromatogram builder were set to 5 minimum consecutive scans, 5e5 minimum absolute height, and 10 ppm *m/z* tolerance. Smoothing was applied. Local minimum features resolver parameters were set to chromatographic threshold 90%, minimum search range retention time 0.05 min, and minimum ratio of peak top/edge of 1.8. Isotope filter with retention time tolerance 0.04 min was used. Join aligner was used for feature alignment, with retention time tolerance set to 0.2. Features not detected in at least 10 samples were removed. <u>Milk and skin</u>: in the chromatogram builder, parameters were set to 4 minimum consecutive scans, 5e3 minimum absolute height, and 10 ppm *m/z* tolerance. Smoothing was applied. In the local minimum features resolver, a chromatographic threshold of 90% was used, with minimum search range retention time of 0.05 min, and minimum ratio of peak top/edge of 1.8. An isotope filter with retention time tolerance 0.04 min was applied. In the join aligner, retention time tolerance was set to 0.2. Features not detected in at least 5 samples were removed. Peak finder, metaCorrelate and ion identity molecular networking were run for all four sample matrices.

For all biofluids, MS/MS spectra (.mgf) with corresponding feature table (.csv) were exported and uploaded to GNPS2^123^ for feature based molecular networking (FBMN)^57^. In short, precursor and fragment ion tolerance was set to 0.02. Networking and annotation parameters were set to cosine similarity > 0.7 and minimum 5 matching peaks. The FBMN jobs are publicly available, see data availability. The network.graphml files from the FBMN jobs were imported to Cytoscape (v 3.10.4)^124^ for visualization and manipulation. MS/MS spectra of unannotated features of interest were run in SIRIUS (v 6.1.1)^125^ and Canopus^55^ was used for chemical class, superclass, and pathway prediction (probability > 0.7). Features of interest were also searched in microbeMASST^33^ and tissueMASST^35^ (February 2026), with parameters set to minimum 4 matching peaks and cosine similarity > 0.7.

### Untargeted Metabolomics Data Analysis

Feature and annotation tables were imported in R 4.4.2 (R Foundation for Statistical Computing, Vienna, Austria) for downstream data analysis and visualization. The annotation tables were merged by feature ID with a priority order; e.g. annotations from the propagated libraries were only used when no prior annotation existed. Further, features annotated as bile acids or *N*-acyl lipids using the propagated bile acid library^60^ or the *N*-acyl lipid library^74^, respectively, underwent post-validation filtering using massQL^126^ (https://massqlpostmn.gnps2.org/)^127^. Predefined queries (bile acid stage 1 and 2, and *N*-acyl lipid queries) were used, with the intensity threshold for bile acids queries modified to 2%. Features annotated using a newly developed carnitine library^56^ also underwent post-validation filtering with the following query:

- *QUERY scaninfo(MS2DATA) WHERE MS2PROD=60.0808:TOLERANCEPPM=20:INTENSITYPERCENT=1 AND MS2PROD=85.0284:TOLERANCEPPM=20:INTENSITYPERCENT=5 AND MS2PROD=144.1019:TOLERANCEPPM=20:INTENSITYPERCENT=1*

For all sample types, data quality was investigated with quality control samples and total extracted peak areas. Blank subtraction was performed, i.e. features detected in the blank samples were excluded unless the peak areas were minimum 5 times that in the samples. For all sample types, features with a retention time (RT) < 0.2 or > 9 min were excluded. Polymers detected in the LC-MS/MS data for fecal samples were removed using the package ‘homologueDiscoverer v 0.0.0.9000’^128^. Additional correlated features identified with the meta correlate function in MZmine^122^ were also excluded. <u>Fecal samples:</u> There was a shift in intensity in the total ion chromatograms (TICs) of the quality controls samples toward the end of the LC-MS/MS run. To correct for this, a ratio between early (<598) and late (>597) QC samples was calculated (ratio = 0.90) and used to adjust peak areas of relevant samples (sample 598-834). Additionally, 3 fecal samples were excluded from the metabolomics analysis due to high TIC or outliers in the PCA. During data analysis, a principal component analysis (PCA) revealed a strong clustering of samples by collection time (**Extended Data Fig. 3a**). This artifact is most likely explained by a change in the kits used for feces collection midway in the study, caused by a lack of availability. To correct for this batch effect, we defined a new categorical variable (Cluster 1 and 2) based on how samples clustered in the PCA, and performed a partial least square discriminant analysis (PLS-DA) using ‘mixOmics v 6.22’ to identify the features driving this clustering (BER = 0.002). A total of 2,795 unannotated features with VIP > 1 were identified and removed, which reduced the batch-related effect and improved clustering (**Extended Data Fig. 3b**). The resulting dataset was used for subsequent downstream analysis. <u>Skin:</u> Similar to feces, there was a clustering by sample collection time in the PCA also for the skin samples (**Extended Data Fig. 3c**). This batch effect was corrected for with the same approach as for fecal samples, with removal of 2,470 unannotated features, and the subsequent dataset used for downstream analysis (**Extended Data Fig. 3d**). Additionally, 13 samples were excluded from downstream analyses due to low or high TIC. <u>Milk:</u> A total of 42 milk samples were excluded due to low internal standard signal (*n*=34) or high TIC (*n*=8). For all sample types, the package ‘caret v 6.0’ was used to remove features with near zero variance. Features were robust center log ratio (RCLR) transformed using ‘vegan v 2.6’ before applying dimensionality reduction techniques. Principal component analysis (PCA) and partial least square discriminant analysis (PLS-DA) models were built using ‘mixOmics v 6.22’^114^. PERMANOVA was used to test centroid separation. The performances of the PLS-DA models were investigated via a 4-folds cross-validation. Variable importance in projection (VIP) scores > 1 were considered significant for group separation unless otherwise is stated. To avoid bias introduced by the antibiotic features themselves, all features annotated as antibiotics were removed before conducting comparative analyses (ABX vs. No ABX/Baseline). For mothers, only metabolic features commonly enriched or depleted after intersecting results from ABX vs Baseline and ABX vs No ABX models, were considered significantly associated with ABX treatment. Log ratios were calculated using the top 100 metabolic features enriched and depleted with ABX. Group differences were assessed with LME including subject id as random effect to account for repeated measures. For milk, the following model was used: *log_ratio ∼ group + maternal_age + bmi_mother + ethnicity + household_income + date_milk_collection + (1|pid)*. For maternal feces and skin, the model was: *log_ratio ∼ group + maternal_age + bmi_mother + ethnicity + household_income + sample_collection_date + (1|pid))*. For infants, downstream analyses were restricted to samples collected during treatment (ABX and No ABX group), as baseline samples were collected on average 4 months prior to treatment. Consistent with the microbiome analysis, only infants indirectly exposed to amoxicillin (with or without clavulanate), cephalexin, penicillin-type drug, and azithromycin and their matching controls were included in differential abundance analyses. Group differences were assessed with LME models including subject id as random effect to account for repeated measures. The following models were used: *log_ratio ∼ group + child_age + delivery_mode + infant_sex + household_income + exclusive_breastmilk + breastfeeding_freq + date_of_skin_collection + (1|pid))* for infant skin and *log_ratio ∼ group + child_age + delivery_mode + infant_sex + type_delivery (preterm/term) + exclusive_breastmilk + breastfeeding_freq + date_of_stool_collection + (1|pid))* for infant feces. In the analyses of the generated log ratio stratified by group and feeding practice, the model was modified accordingly: *log_ratio ∼ feeding_group + child_age + delivery_mode + infant_sex + type_delivery (preterm/term) + date_of_stool_collection + (1|pid)).* For comparisons by source (mother vs infant), the model was: *log_ratio ∼ source + sample_timepoint + (1|subject_id))* for both feces and skin. ‘Emmeans’ (v 2.0.1) was used to compute estimated marginal means from the LME models and pairwise *p* values using Tukey for adjustment. The volcano plot was generated by log2-transforming peak areas and then using LME to calculate β-estimates and p values. Models were adjusted for maternal age, BMI, ethnicity, household income, and sample collection date, as well as repeated measures by including subject id as a random effect. β-estimates and p values for “group” were extracted using ‘broom.mixed’^129^ and corrected for multiple testing using the BH correction. Adjusted p values were then -log10 transformed. The UpSet plot was generated using ‘UpSetR v 1.4’^130^. Only features with mean VIP>1.5 were included in the visualization. For analyses assessing detection of antibiotics and their metabolites or structural analogs, a stringent peak area threshold (skin >30,000; feces > 10,000) was applied to limit false positive detections. Additionally, a feature was only considered a true detection if the annotated antibiotic was concordant with the antibiotic reported in the corresponding metadata.

## Data Availability

Metagenomics sequencing data is available on EBI/ENA under the accession code <u>PRJEB108678</u>. Untargeted LC-MS/MS data for this study are publicly available at GNPS/MassIVE (https://massive.ucsd.edu/) under the accession codes: <u>MSV000099924</u> (feces), <u>MSV000100756</u> (skin), and <u>MSV000100755</u> (human milk). Associated FBMN jobs are publicly available at GNPS2:

- Feces: https://gnps2.org/status?task=aa40dc3f95fb436a9c41e03acb76a67f (default and propagated bile acid library), https://gnps2.org/status?task=7cced9aed86e49928eb6a9722f1aa44e (drug library), https://gnps2.org/status?task=117fe60f9ddd4af6bd7a0df69e95d98f (carnitine library), https://gnps2.org/status?task=6dcae285e2b7430ba7d768dffc130c17 (synthesis library, filtered).
- Milk: https://gnps2.org/status?task=955fc80bd2b749518ce06371e58e8444 (default and propagated bile acid library), https://gnps2.org/status?task=a0de3b4a62cf4152ac2e700c0df46dcb (drug library), https://gnps2.org/status?task=a8c42c37b87246dc9bc619225f129dbb (carnitine library), https://gnps2.org/status?task=577a8d3d5fdc4de78b18efd7b73d2c52 (synthesis, filtered).
- Skin: https://gnps2.org/status?task=13b7b6dc7e1d4ea691fcbedc5ae8e08b (default and propagated bile acid library),

https://gnps2.org/status?task=92984171f4b44cf180459f7fd9376333 (drug library), https://gnps2.org/status?task=e6f5e0db29a84be9a360c1a731af7f42 (carnitine library), https://gnps2.org/status?task=51090b15ed994b5285121d7d298d4f0f (synthesis, filtered).

## Code Availability

Code used for metabolomics analysis and to generate figures is available at https://github.com/kinekvitne/manuscript_MPRINT. Code used for metagenomics and multi-omics analyses and to generate figures is available at https://github.com/simonezuffa/Manuscript_MPRINT_Clinical

